# Small terrestrial mammal distributions in Simien Mountains National Park, Ethiopia: A reassessment after 88 years

**DOI:** 10.1101/771428

**Authors:** Evan W. Craig, William T. Stanley, Julian C. Kerbis Peterhans, Josef Bryja, Yonas Meheretu

## Abstract

Little is known about the distribution and ecology of small mammals inhabiting Simien Mountains National Park despite the presence of mostly endemic species. Prior to this study, the most comprehensive dataset was collected in 1927. This provides a unique opportunity to assess the possible role of climate change over the last 88 years on the elevational distribution of mammals in the Ethiopian highlands. Between September and November 2015, three of us (EWC, WTS, YM) collected non-volant small mammals at four sites (2900, 3250, 3600, and 4000 m a.s.l.) along the western slope of the Simien Mountains using standardized sampling. Over a four-week period we recorded 13 species, comprising 11 rodents and two shrews, all endemic to the Ethiopian Plateau. We found greatest species richness at mid-elevations (3250 m), consistent with a general pattern found on many other mountains worldwide but less so in Africa. We discovered one potentially new species of shrew. No previously unrecorded rodent species were observed. Finally, we compared our species distribution results to the 1927 dataset and found upward elevational shifts in species ranges, suggesting the role and influence of climate change on the small mammal community. Simien Mountains National Park represents an exceptionally valuable core area of endemism and the best protected natural habitat in northern Ethiopia.

---

Understanding the distributions of organisms along elevational gradients is vital to comprehend the evolution and ecology of montane biotic systems, and to facilitate effective conservation strategies to maintain them. Gathering such baseline data is necessary to monitor any changes that may be occurring due to climate change, ecological perturbations, and impacts caused by human activity (Walther et al. 2002; Rowe et al. 2010; Chen et al. 2011; Sundqvist et al. 2013). This has been illustrated by Moritz et al. (2008) who revealed significant elevational shifts in the distributions of various mammalian species in Yosemite National Park, USA over a 100 year period as a result of increasing global temperatures, and by Ejigu et al. (2017) who associated human-related activities with a shift in walia ibex (*Capra walie*) ranges in the Simien Mountains of Ethiopia.

Temporal variations in the elevational distributions of natural communities provide an indication to the rate at which change occurs. Accurately predicting the ecological consequences of changing habitats, however, requires an understanding of the relationship of biotic variables with species occurrence. One notable development recently changed what had been a fundamental assumption regarding biodiversity along elevational gradients. Emerging evidence suggests that species richness is typically greatest at mid-elevations (Rahbek 2005), as opposed to the prior notion of an inverse linear relationship between richness and elevation (MacArthur 1972). Support for small mammals exhibiting this mid-elevational peak was provided in 2005 by the assessment of 56 published montane diversity surveys, only four of which did not report greatest richness at mid-elevations (McCain 2005). However, despite mounting global evidence in favor of this general pattern, such “hump-shaped” distributions of non-volant small mammals in Africa have been infrequently documented (Taylor et al. 2015).

Ethiopia is home to the largest Afroalpine ecosystem (areas above approximately 3200 m), which is disjointedly distributed across several isolated massifs (Yalden 1988). The isolated massifs are the result of the great Ethiopian volcanic eruptions ca. 30 Ma (Hofmann et al. 1997). High isolated plateaus combined with unique environmental conditions have resulted in significant faunal and floral endemism. With the exception of two murid rodents, nearly all (96%) of the 55 mammals currently reported as endemic to Ethiopia are restricted to the plateau, and approximately half (47%; *n* = 26) live in the highlands (Lavrenchenko and Bekele 2017). The IUCN Small Mammal Specialist Group has identified these Ethiopian montane grasslands and woodlands as a key region for conservation (IUCN 2019).

Simien Mountains National Park (SMNP) was established in 1969 and recognized as a UNESCO World Heritage Site in 1978 (UNESCO 2019). Home to Ras Dashen, the highest mountain in Ethiopia and tenth highest in Africa, the park has long been a subject of interest to researchers with its populations of rare and endemic species. This montane “sky island” contains iconic mammalian species, such as the Ethiopian wolf (*Canis simiensis*), gelada (*Theropithecus gelada*), and walia ibex (*Capra walie);* the smaller mammals are less well known.

In 2015, we (EWC, WTS, YM) conducted an elevational survey in SMNP to reassess the small mammal community. This collaborative study between Mekelle University and the Field Museum of Natural History (FMNH) marked nearly a century-long return to the Simien Mountains following former FMNH Curator of Zoology, W.H. Osgood’s historic Chicago Daily News Abyssinian Expedition (hereafter “Abyssinian Expedition”) of 1927. Although Osgood published some results of the Abyssinian Expedition, including species accounts and descriptions of new species (Fuertes and Osgood 1936; Osgood 1936), sampling results such as species abundance and distribution data were not reported. By referencing Osgood’s collection and associated documents from the Abyssinian Expedition deposited at FMNH, we provide an 88-year reassessment of small mammal distributions along the same route in SMNP.

Outside Africa, small mammal redone surveys have revealed elevational shifts in species ranges, likely as a response to climate change (Moritz et al. 2008; Rowe et al. 2010). Given the global influence of climate change, we hypothesize that the SMNP community has responded similarly to warming temperatures through elevational shifts in species distributions. Our study objectives were: 1) to assess the current elevational distribution of non-volant small mammal diversity in Simien Mountains National Park, Ethiopia; 2) to evaluate the efficacy of, and identify sampling biases associated with different trapping techniques for small mammal diversity assessments; and 3) to compare our results to those of the Abyssinian Expedition in 1927, and assess shifts in species distributions.

## Materials and methods

### Study area

Simien Mountains National Park covers an area of 412 km^2^ and is located between about 13°-13.5° N, and 37.8°-38.5° E in the Amhara Regional State, Ethiopia (Fig. 1). The mountain range is entirely made up of flood basalt, the product of volcanic eruptions during the Oligocene-Miocene epoch (Hofmann et al. 1997). The landscape within SMNP exhibits a variety of habitat types along its elevational profile that range from Afromontane forests at the base of the massif to Afroalpline meadows nearest the summit. The Simien Mountains experience a unimodal pattern of rainfall that varies in volume from north (drier) to south (wetter) due to the 1000 m-high escarpment’s rain shadow effect (Jacob et al. 2017). The wet season generally occurs between May and September. Rainfall data collected at Chennek camp (3600 m.a.s.l. site sampled in this study) shows an annual total average of 825.7 mm with the highest monthly averages occurring in July (287.2 mm; Chernet 2015).

**Fig. 1.**
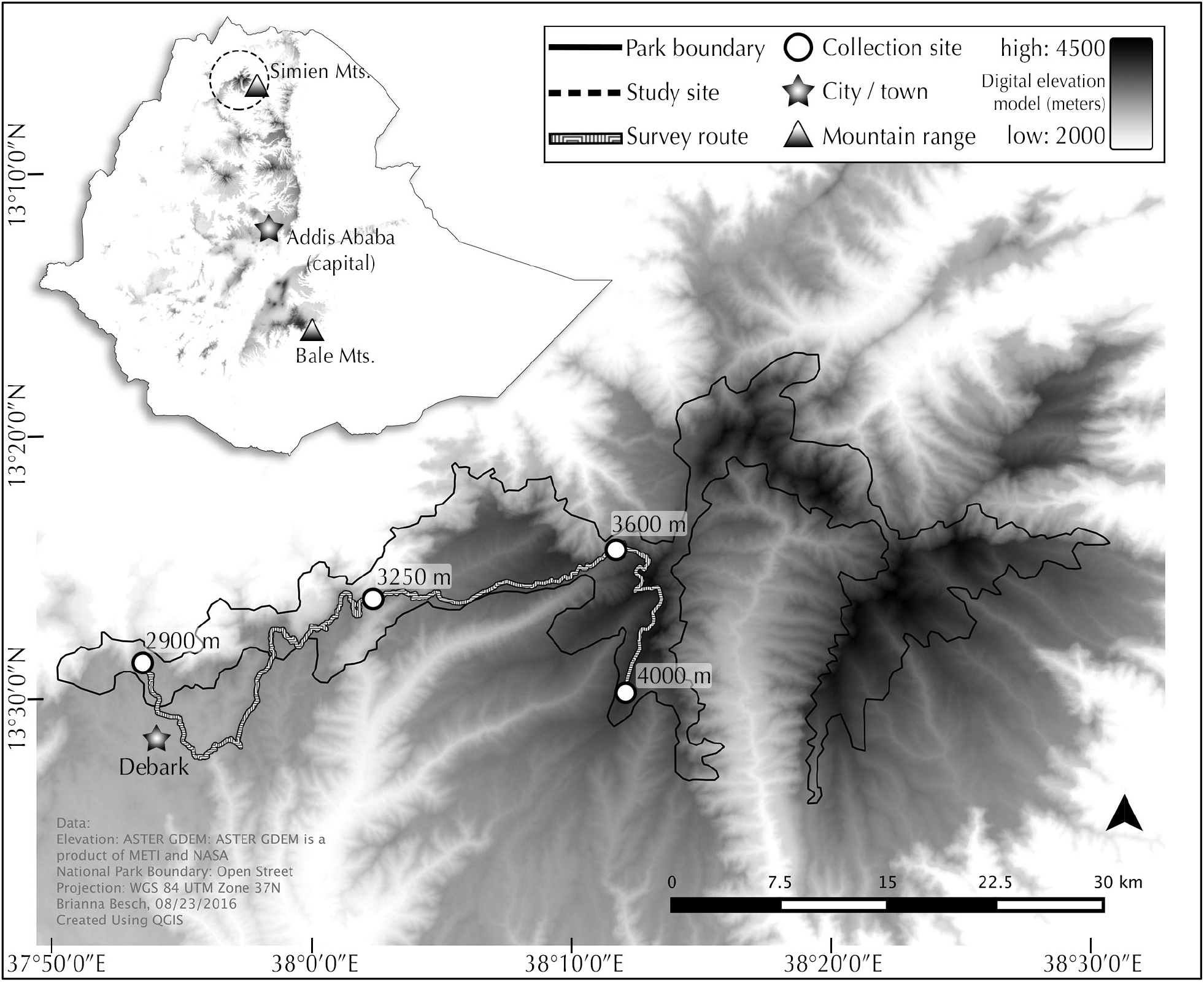
Map of Simien Mountains National Park, Ethiopia showing study sites, survey route, and country reference.

A single unpaved road provides vehicle access to settlements and scout camps along the western aspect of the massif. Between 21 September and 28 November 2015, we sampled the small mammal populations along this route at four different sites spanning an elevational range of roughly 1100 m (Fig. 2). Elevations, camp names, specific localities, sampling dates, and habitat notes associated with each site are presented below. Elevations for each site reference the center of the associated camp (to the nearest 50 m) with sampling efforts extending roughly 50 m above or below this point.

**Fig. 2.**
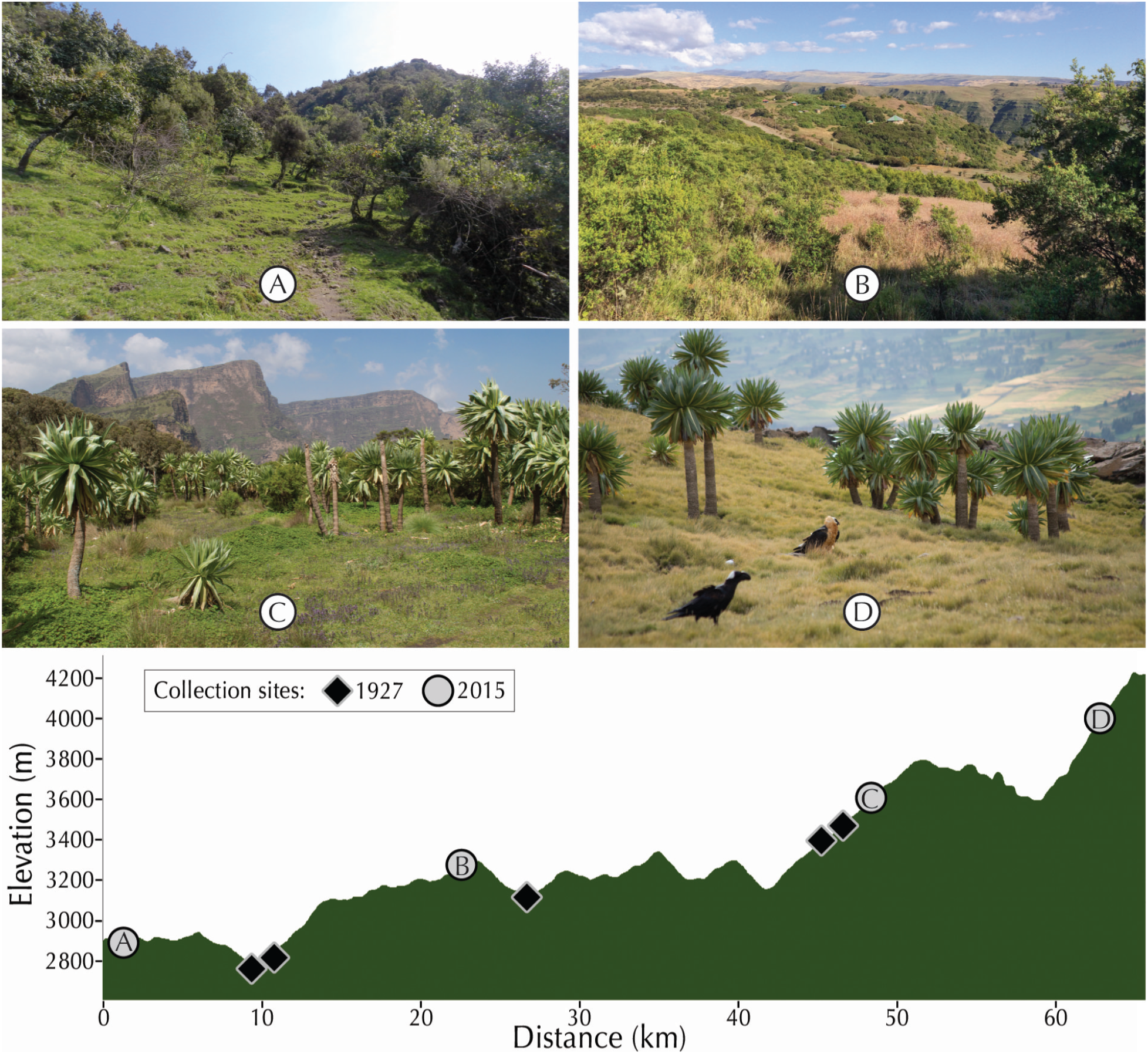
Typical habitats photographed at each site sampled between September and November 2015 in Simien Mountains National Park, Ethiopia. Site elevations and associated camp names are: (A) 2900 m, Lima Limo; (B) 3250 m, Sankaber; (C) 3600 m, Chennek; and (D) 4000 m, Sabat Minch. Elevational profile (bottom) of survey route showing approximate locations sampled in 1927 by the Chicago Daily News Abyssinian Expedition, and by us in 2015 (letters correspond to habitat photos). Photographs by YM (A), and EWC (B-D).

Site 1: 2900 m; Lima Limo; 13.19°N, 37.89°E; 5-10 October, 2015.—The lowest site was positioned within Afromontane forest where the dominant tree species consisted of *Juniperus procera, Olea europaea, Rapanea simensis*, and *Hagenia abyssinica*. This site also included isolated patches of *J. procera* and eucalyptus (*Eucalyptus* sp.) monocultures.

Site 2: 3250 m; Sankaber; 13.23°N, 38.04°E; 22-28 November, 2015.—This site was located in a transition zone between Afromontane forest and ericaceous heathland where *Hypericum revolutum* was prevalent. *Nuxia congesta* was established on the steepest slopes and escarpments.

Site 3: 3600 m; Chennek; 13.26°N, 38.19°E; 21-27 September, 2015.—At this site was ericaceous heathland. Giant heather (*Erica arboea*) and the endemic giant lobelia (*Lobelia rhynchopetalum*) dominated the landscape.

Site 4: 4000 m; Sabat Minch; 13.17°N, 38.20°E; 28 September-4 October, 2015.—At the highest site *E. arborea* was scarce, while *L. rhynchopetalum* remained abundant among the grassy Afroalpine meadows.

### Sampling protocol

Trapping techniques used to capture non-volant small mammals consisted of pitfall buckets, Sherman live traps, and snap traps. We used this combination of traps to maximize our potential for sampling the highest diversity of mammals possible, as capture probabilities for each type can vary significantly among species. Pitfall lines consisted of 11 plastic buckets (depth = 29.5 cm, rim diameter = 28.5 cm) spaced 5 m apart, installed in a linear sequence so that the upper rim of the opening was flush with the ground. Pitfall lines were then fitted with a 50 cm high plastic fence bisecting the opening of each bucket and running the length of the line. Trap lines consisted of 50 traps, comprised of 14 medium-sized Sherman Traps 23 × 9.5 × 8 cm (H.B. Sherman Traps Inc., Tallahassee, Florida, USA) and various snap traps: 14 Museum Specials, 14 × 7 cm; 16 Victor Rat Traps, 17.5 × 8.5 cm (both manufactured by Woodstream Corporation, Lititz, Pennsylvania, USA); and 6 small snap traps manufactured in the Czech Republic that we called the “Czech Mouse Trap”, 9.5 × 4.5 cm. Selection of trap type and placement of traps and trap lines at each site was based on collector discretion, influenced primarily by habitat and microhabitat features. Distances between traps within trap lines were not consistent though never exceeding 10 m. Additional details on placement of traps are provided in Stanley et al. (2011). Sites received five to seven consecutive days of trapping (detailed sampling efforts for each site are provided in Table 2). Traps and buckets were checked every morning (~0700 hrs) and afternoon (~1800 hrs). All traps (but not buckets) were baited and refreshed each afternoon with a mixture of peanut butter and canned tuna. For each 24-hour period we utilized a digital thermometer to record the minimum and maximum temperatures, and a rainfall gauge to measure any precipitation (Supplementary Data SD1).

### Species identification

Liver or spleen tissue samples were stored in 96% ethanol or Dimethyl sulfoxide (DMSO) until DNA extraction. The complete mitochondrial genome for cytochrome *b* was amplified and Sanger-sequenced (using the protocol described in Bryja et al. 2014) in select individuals, representing different morphotypes (= species) and elevations. Obtained sequences were aligned with both published, and our own unpublished data for the confirmation of species identification. This was done by producing maximum likelihood tree by FastTree (Price et al. 2009) for each genus, and visual exploration of phylogenetic affinities of specimens from Simien Mts. (for specific references to each genus, see Table 1). All new sequences were submitted to GenBank under accession numbers MN223586-MN223667 (see Supplementary Data SD2).

**Table 1.** Elevational distribution of small mammals in Simien Mountains National Park, September-November 2015. Numbers in parentheses indicate how many individuals were used for genetic identification, based on phylogenetic analysis of cytochrome *b* gene and comparison with published data (specified in Reference column). GenBank accession numbers and museum (FMNH) numbers of genotyped vouchers are provided in Supplementary Data SD2.

| Elevation | 2900 m | 3250 m | 3600 m | 4000 m | Totals | Reference |
| --- | --- | --- | --- | --- | --- | --- |
| Shrew species |  |  |  |  |  |  |
| *Crocidura baileyi* | 0 | 14 (3) | 53 (8) | 22 (4) | 89 | (Lavrenchenko et al. 2009) |
| *Crocidura* sp. indet. | 8 (2) | 18 (4) | 8 (3) | 0 | 34 | (Lavrenchenko et al. 2009) |
| Rodent species |  |  |  |  |  |  |
| *Arvicanthis abyssinicus* | 0 | 0 | 32 (3) | 53 (4) | 85 | (Bryja et al. 2019) |
| *Dendromus lovati* | 0 | 3 (3) | 8 (4) | 6 (3) | 17 | (Lavrenchenko et al. 2017) |
| *Dendromus mystacalis* | 0 | 10 (1) | 0 | 0 | 10 | (Lavrenchenko et al. 2017) |
| *Desmomys harringtoni* | 0 | 2 (2) | 0 | 0 | 2 | (Bryja et al. 2017) |
| *Lophuromys simensis* | 1 (1) | 61 (4) | 19 (4) | 0 | 81 | (Lavrenchenko et al. 2004) |
| *Mus mahomet* | 0 | 7 (2) | 0 | 0 | 7 | (Bryja et al. 2014) |
| *Mus imberbis* | 0 | 3 (3) | 1 (1) | 0 | 4 | (Bryja et al. 2014) |
| *Otomys simiensis* | 0 | 14 (3) | 0 | 0 | 14 | (Taylor et al. 2011) |
| *Otomys typus* | 0 | 0 | 27 (4) | 8 (3) | 35 | (Taylor et al. 2011) |
| *Stenocephalemys albipes* | 8 (4) | 19 (5) | 0 | 0 | 27 | (Bryja et al. 2018) |
| *Stenocephalemys* sp. “A” | 0 | 0 | 19 (4) | 48 (7) | 67 | (Bryja et al. 2018) |
| Total # shrew individuals | 8 | 32 | 61 | 22 | 123 |  |
| Total # rodent individuals | 9 | 119 | 106 | 115 | 349 |  |
| Total # shrew species | 1 | 2 | 2 | 1 | 2 |  |
| Total # rodent species | 2 | 8 | 6 | 4 | 11 |  |

### Statistical analysis

We use “trap-night”, “bucket-night”, and “sample-night” to clearly enumerate the sampling effort. The term “trap-night”/“bucket-night” is defined as one set trap/bucket for a 24-hour period, in this case 0700 to 0700 hours. “Sample-night” refers to the combined sampling effort (of both traps and buckets) for the same period. We describe the success rate of each method in terms of “trap success” and “bucket success”, while “sample success” refers to the combined success rate of both traps and buckets. The success rate for each technique is calculated by dividing the number of individuals caught in traps or buckets by the number of trap-nights or bucket-nights, and multiplying by 100. Sample success is calculated by dividing the overall number of individuals captured from both methods by the number of sample-nights. Additional details are provided in Stanley et al. (2014).

To assess the sufficiency of our sampling efforts, we mapped species accumulation curves and total capture abundances for each night of trapping at each site. The Shannon index (*H*) was used to measure species diversity,

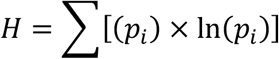

Where *p_i_* is the proportion of the total sample at each site represented by species *i*. Evenness (*E_H_*), was then calculated by dividing *H* for a given site from its maximum possible diversity (*H_max_*=*lnS*), where *S* is the number of species observed. All statistical analyses were performed using Microsoft Excel.

### Ethics statement

Permission for the collection and export of specimens was provided by the Federal Democratic Republic of Ethiopia, Ethiopian Wildlife Conservation Authority No. 229/27/08. Approval for the import of specimens into the USA was provided by the US Fish and Wildlife Service (3177-1/11/2016). All euthanized specimens followed the protocol approved by the American Society of Mammalogists (Sikes et al. 2011). The study was approved by the Field Museum of Natural History Institutional Animal Care and Use Committee (09-3).

### Abyssinian Expedition data collection

To confirm species identities and map small mammal distributions in the Simien Mountains in 1927 we referenced maps, specimen records (i.e. field catalog), journal entries, and voucher specimens deposited by the Abyssinian Expedition at FMNH. Maps provided an overview of the survey route and timeline. The field catalog provided locality and elevation data for each specimen record. Journal entries often contained contextual or direct references to camps and landmarks that were used to corroborate locality data. Morphological analyses of voucher specimens confirmed species identities. Original species identities were made by W.H. Osgood.

### Survey comparison and analysis

From March 15 to April 4, 1927, the Abyssinian Expedition sampled along the western aspect of the Simien Mountains, including many localities within what is now SMNP. To maximize our ability to make direct comparisons between the two surveys, we included only localities sampled in 1927 located within the present-day boundary of SMNP in our assessment. A total of six localities met this criteria at the following elevations (rounded to the nearest 50 m): 2700 m, 2750 m, 2800 m, 3050 m, 3350 m, and 3400 m (Fig. 2). For the same reason, we consolidated localities from both surveys into four elevational ranges based on present dominant vegetation belts in SMNP: Afromontane forest (2000-2900 m), Afromontane forest/ericaceous heathland (2900-3300 m), ericaceous heathland (3300-3700 m), and Afroalpine meadows (3700+ m; Jacob et al. 2017). For clarity, we use the term “elevation zones” when referencing these ranges in the text.

## Results

We captured a total of 472 small mammals (349 rodents and 123 shrews) during our survey (Table 1). The rodents were represented by seven genera comprising 11 species, and the shrews by a single genus (*Crocidura*) with two species. Our total sampling effort was an accumulated 6,273 sample-nights (Table 2). In 4,700 trap-nights we captured 380 small mammals with a trap success of 8.1%, 319 were rodents (6.8% trap success), and 61 were shrews (1.3% trap success). The pitfall effort of 1,573 bucket-nights yielded 92 small mammals with an overall bucket success of 5.8%, 30 were rodents (1.9% bucket success), and 62 were shrews (3.9% bucket success). *Crocidura baileyi* (weight; 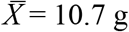) was readily caught by various traps in all three of the elevations in which it was recorded. The potentially new species *Crocidura* sp. indet. (weight, 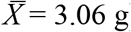) was only captured by pitfalls. The majority of rodents were captured by traps, although not an uncommon occurrence in pitfall buckets. Of the 7 rodent species found in buckets (*Dendromus lovati, Dendromus mystacalis, Lophuromys simensis, Mus imberbis, Otomys simiensis, Otomys typus, and Stenocephalemys* sp. “A”), only *D. mystacalis* (weight; 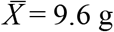) was absent from traps.

**Table 2.** Trapping results of rodents and shrews in Simien Mountains National Park, September-November 2015.

| Elevation | 2900 m | 3250 m | 3600 m | 4000 m | Totals |
| --- | --- | --- | --- | --- | --- |
| Buckets |  |  |  |  |  |
| # bucket-nights | 330 | 451 | 396 | 396 | 1573 |
| # individuals | 9 | 43 | 26 | 14 | 92 |
| % bucket success | 2.7 | 9.5 | 6.6 | 3.5 | 5.8 |
| # species | 2 | 6 | 3 | 4 | 10 |
| # shrews | 8 | 27 | 21 | 6 | 62 |
| % bucket success - shrews | 2.4 | 6 | 5.3 | 1.5 | 3.9 |
| # shrew species | 1 | 2 | 2 | 1 | 2 |
| # rodents | 1 | 16 | 5 | 8 | 30 |
| % bucket success - rodents | 0.3 | 3.5 | 1.3 | 2 | 1.9 |
| # rodent species | 1 | 4 | 1 | 3 | 7 |
| Traps |  |  |  |  |  |
| # trap-nights | 1000 | 1300 | 1200 | 1200 | 4700 |
| # individuals | 8 | 108 | 141 | 123 | 380 |
| % trap success | 0.8 | 8.3 | 11.8 | 10.3 | 8.1 |
| # species | 1 | 7 | 7 | 5 | 8 |
| # rodents | 8 | 103 | 101 | 107 | 319 |
| % trap success - rodents | 0.8 | 7.9 | 8.4 | 8.9 | 6.8 |
| # rodent species | 1 | 6 | 6 | 4 | 7 |
| # shrews | 0 | 5 | 40 | 16 | 61 |
| % trap success - shrews | 0 | 0.4 | 3.3 | 1.3 | 1.3 |
| # shrew species | 0 | 1 | 1 | 1 | 1 |
| Totals |  |  |  |  |  |
| # sample-nights | 1330 | 1751 | 1596 | 1596 | 6273 |
| % sample success - shrews | 0.6 | 1.8 | 3.8 | 1.4 | 2 |
| % sample success - rodents | 0.7 | 6.8 | 6.6 | 7.2 | 5.6 |
| % sample success - overall | 1.3 | 8.6 | 10.5 | 8.6 | 7.5 |

Rainfall was positively correlated with sampling success for shrews but not rodents during the survey. At 2900 m the correlation was significant (*p* = 0.049) and most pronounced, with an increase in shrew captures coinciding with the period of heaviest rainfall. Product-moment correlation coefficients (*r*) for rainfall and shrew capture success (buckets and traps combined) are 0.88 (2900 m), 0.67 (3600 m), 0.17 (4000 m), and for rodents are 0.11 (2900 m), 0.39 (3600 m), and 0.24 (4000 m; Supplementary Data SD3).

The number of small mammals captured ranged from a low of 17 (1.3%) individuals at 2900 m, to 167 (10.5%) individuals at 3600 m (Table 2). The number of shrew captures by site share the same lower and upper limits, with eight (0.6%) at 2900 m, and 61 (3.8%) at 3600 m. The number of rodents captured by site ranges from nine (0.7%) at 2900 m to 119 individuals (6.8%) at 3250 m. Species richness was greatest at 3250 m (10 species). This elevation also contained all four species (*D. mystacalis, Desmomys harringtoni, M. mahomet*, and *O. simiensis*) that occurred exclusively at one site along the survey transect (Table 1). Diversity was greatest at 3250 m (Shannon index, *H* = 1.86). Values for evenness were similar across elevations, though highest at 3600 m (*E_H_* = 0.86; Fig. 3).

**Fig. 3.**
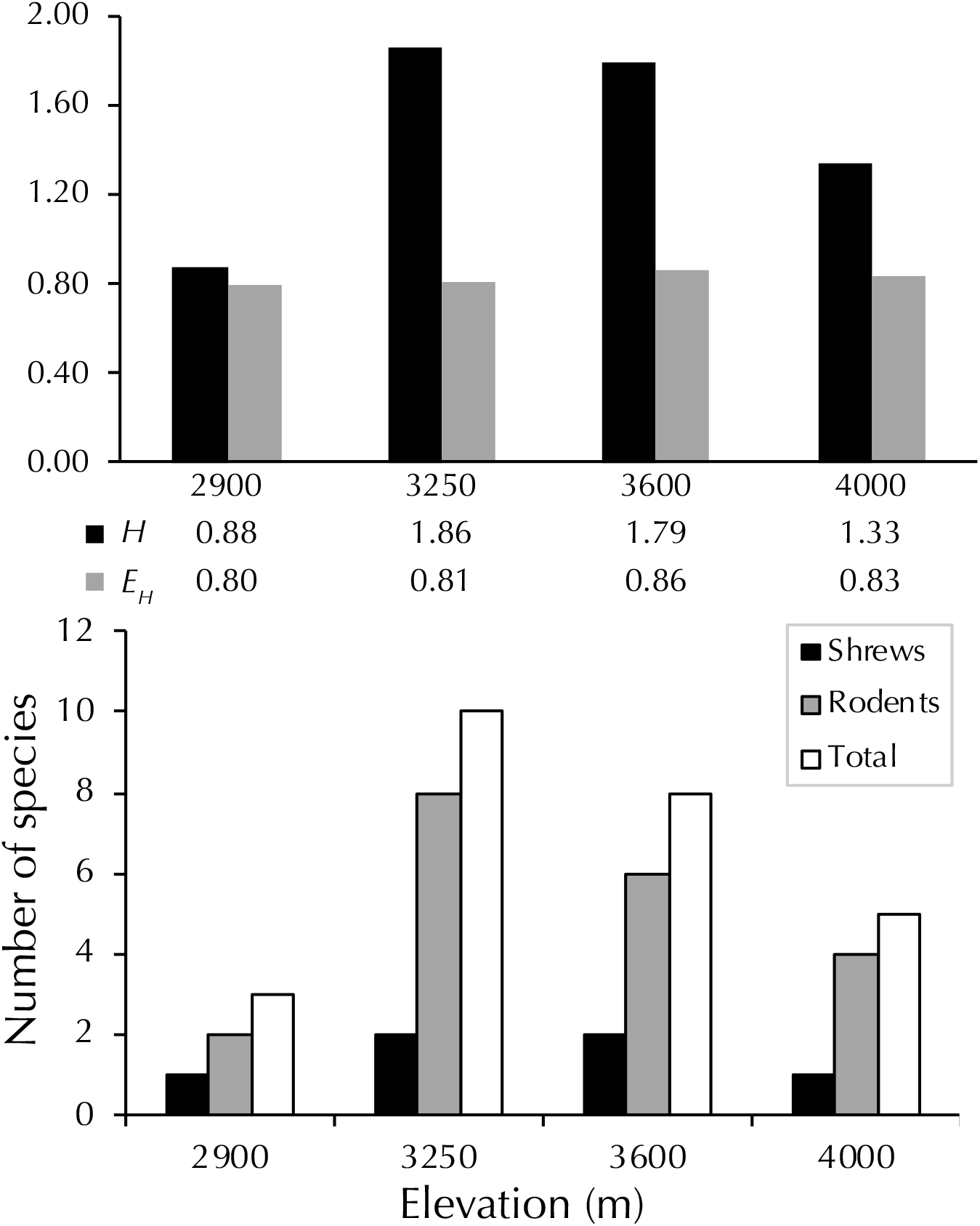
Diversity of small mammals at different elevations in Simien Mountains National Park. Species richness (bottom), diversity, and evenness (rodents and shrews combined; top) are presented for each site elevation. Calculations for Shannon’s diversity (H) and evenness (E_H_) are provided in the text.

The total number of species recorded at each site was reached by the second day of trapping except at 2900 m, where *L. simensis* was recorded on the fourth day. Thus, species accumulation curves reached an asymptote for each site. (Fig. 4).

**Fig. 4.**
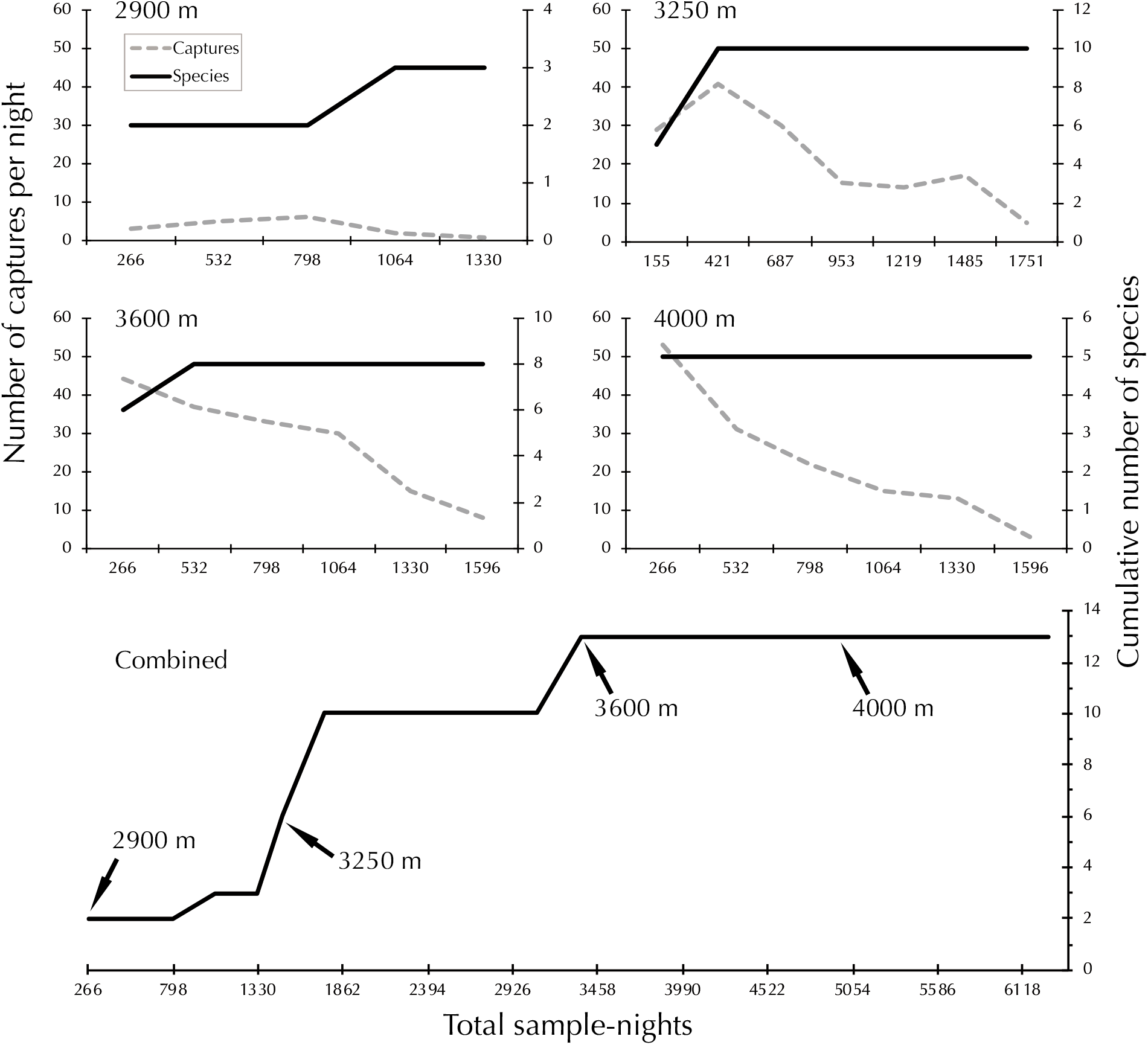
Species accumulation curves for each site (top). The dashed line represents the number of individuals captured. The solid line represents the cumulative number of new species recorded. Species accumulation for all sites combined (bottom); arrows indicate initiation of trapping effort at each site.

### Abyssinian Expedition

A total of 101 small mammals consisting of nine species (1 shrew, and 8 rodents) were recorded in the Simien Mountains by the Abyssinian Expedition of 1927 (Table 3). No terrestrial small mammal species were recorded that were absent from our 2015 survey. Conversely, we collected one shrew (*Crocidura* sp. indet.) and three rodent taxa (*D. harringtoni, D. mystacalis*, and *O. simiensis*) not reported by the 1927 expedition. Of these, *D. harringtoni, D. mystacalis*, and *O. simiensis* were captured exclusively at the 3250 m site in 2015. Both surveys sampled the Afromontane forest, Afromontane forest/ericaceous heathland, and ericaceous heathland elevation zones (2000-2900 m, 2900-3300 m, and 3300-3700 m, respectively) while only the 2015 survey reached the Afroalpine meadows (3700+ m).

**Table 3.** Number of individuals captured for each species by elevation zone in 1927 and 2015 (in parentheses) in Simien Mountains National Park. Dashes indicate no recorded individuals. Species not recorded by the Abyssinian Expedition in 1927 are indented. Elevation zone abbreviations: Afromontane forest (AMF); Afromontane forest/ericaceous heathland belt (AMF/EH); ericaceous heathland (EH); Afroalpine meadow (AAM).

| Species | Elevation zones |  |  |  |
| --- | --- | --- | --- | --- |
|  | AMF | AMF/EH | EH | AAM<sup>a</sup> |
|  | 2000-2900 m | 2900-3300 m | 3300-3700 m | 3700+ m |
| *Crocidura baileyi* | - (-) | 2 (14) | 3 (53) | (22) |
| *Crocidura* sp. indet. | - (8) | - (18) | - (8) | (-) |
| *Arvicanthis abyssinicus* | 6 (-) | 10 (-) | 2 (32) | (53) |
| *Dendromus lovati* | 1 (-) | 1 (3) | 3 (8) | (6) |
| *Dendromus mystacalis* | - (-) | - (10) | - (-) | (-) |
| *Desmomys harringtoni* | - (-) | - (2) | - (-) | (-) |
| *Lophuromys simensis*<sup>b</sup> | 8 (1) | 6 (61) | 12 (19) | (-) |
| *Mus imberbis*<sup>c</sup> | 2 (-) | - (3) | 1 (1) | (-) |
| *Mus mahomet* | 1 (-) | - (7) | - (-) | (-) |
| *Otomys simiensis* | - (-) | - (14) | - (-) | (-) |
| *Otomys typus* | 4 (-) | 1 (-) | 10 (27) | (8) |
| *Stenocephalemys albipes*<sup>d</sup> | 13 (8) | - (19) | - (-) | (-) |
| *Stenocephalemys* sp. "A"<sup>e</sup> | 5 (-) | 7 (-) | 3 (19) | (48) |
| Totals | 40 (9) | 27 (107) | 34 (159) | (137) |

<sup>a</sup>Not sampled in 1927, 2015 results are provided for context. Originally named by Osgood in 1927:
<sup>b</sup>*Lophuromys flavopunctatus simensis*, <sup>c</sup>*Muriculus imberbis imberbis*, <sup>d</sup>*Myomys albipes*, and <sup>e</sup>*Stenocephalemys griseicauda*.

Among the taxa collected in both expeditions, six species (*Arvicanthis abyssinicus*, *Dendromus lovati, M. imberbis, Mus mahomet, O. typus*, and *Stenocephalemys* sp. “A”) were recorded at lower elevation zones in 1927 than they were in 2015. Conversely, no species were recorded at higher elevation zones in 1927 than they were in 2015. Details pertaining to field methodology were mostly undocumented by the Abyssinian Expedition, including sampling effort and trapping procedures. As a result, relative abundance data could not be generated.

## Discussion

### Community reassessment: possible elevational shifts over 88 years

Small mammal sampling along the same route in the Simien Mountains after almost nine decades provides a unique opportunity to analyze the changes in the distribution of species. Our results indicate that multiple species have experienced some form of upward movement in their elevational ranges since 1927 (Table 3). For example, the Abyssinian Expedition collected *A. abyssinicus* across all elevation zones sampled. Despite the majority of these individuals (89%; *n* = 16) being captured from the lowest elevation zones (Afromontane forests and Afromontane/ericaceous heathland belt; 2000-3300 m) in 1927, we did not collect any in this range. We began collecting *A. abyssinicus* in ericaceous heathlands (3300-3700 m; *n* = 32) and found them in even greater abundance amongst the higher Afroalpine meadows (3700+ m; *n* = 53). We found a similar pattern for *O. typus* and *Stenocephalemys* sp. “A”; both recorded across all three elevation zones sampled by the 1927 Abyssinian Expedition. *Otomys typus* was absent from our trapping in the Afromontane forest and Afromontane forest/ericaceous heathland zones (2000-3300 m) but was found higher in the ericaceous heathland (3300-3700 m; *n* = 27) and Afroalpine meadows (3700+ m; *n* = 8). We also found *S*. sp. “A” only in these two highest elevation zones where it exhibited a notable increase in abundance amongst the high Afroalpine meadows (3700+ m; *n* = 48), more than doubling in yield from the lower ericaceous heathlands (3300-3700; *n* = 19). The distributional shift in *S. albipes* is again in the same direction. In 1927 it was found only in the lowest Afromontane forest, while our recent survey found it abundant also in the Afromontane forest/ericaceous heathland zone at 3250 m. Other species were less abundant, and it is therefore difficult to make strong conclusions, but they also suggest similar distributional changes. In 2015, *D. lovati* was not captured in the Afromontane forest (2900 m), but it was documented in this zone in 1927. Similarly, the lone specimen of *M. mahomet* documented by the Abyssinian Expedition was collected in the Afromontane forest (2000-2900 m), while we collected all individuals (n = 7) in the higher Afromontane forest/ericaceous heathland zone (3000-3300 m).

Considering our intensive sampling, the absence of multiple species at lower elevations in 2015—despite having been documented there in 1927—indicates upward range contractions have occurred. No species experienced a downward range contraction, as we collected all species at the highest elevation zones from which they were documented in 1927. The most parsimonious explanation for these changes include elevational shifts in habitat types, caused by a changing climate. It is expected that increasing mean temperatures associated with global warming might shift particular ecosystems to higher latitudes and elevations (Sundqvist et al. 2013). As most small mammal species are habitat specialists, one would also expect similar shifts in their distribution as we see in our data. An indication of this climatic effect occurring in SMNP is a rising treeline in areas receiving low anthropogenic pressure (Jacob et al. 2017). However, there are also other factors that should be considered when comparing our results with those obtained in 1927. First, the degradation of habitats by human activities (e.g. overgrazing, deforestation) has intensified in last decades, especially at lower elevations. It is therefore possible that upward elevational shifts were forced by human activities. Second, we have not genotyped the material from 1927 expedition and there is a possibility that material of two genera collected at lower elevations belong to other species than those we observed in higher elevation zones. Lower elevations of northern Ethiopia can be inhabited by *Arvicanthis niloticus*, but it has not been yet reported from SMNP despite intensive sampling in the last decade (Bryja et al. 2019). Also, we have not performed any genetic analysis of *Otomys* collected in 1927. However, our morphological analysis of *Otomys* collected in 1927 confirmed their belonging to the *O. typus* morphotype (sensu Taylor et al. 2011). Third, our recent survey employed pitfall buckets that allowed us to collect some taxa difficult to document by standard traps (e.g. *Dendromus, Mus, Crocidura*). In particular, the absence of species with low body mass can be attributed to the fact that the Abyssinian Expedition employed only traps, while we collected most of these specimens using buckets. Finally, all four species absent in 1927 appear to be more or less specialized to the band of transition between Afromontane forest and ericaceous heathland. This habitat corresponds to the 3250 m site (Sankaber Camp) in our survey and is where we found *C*. sp. indet. in greatest abundance, and *Dendromus mystacalis, Desmomys harringtoni*, and *O. simiensis*, exclusively. During the survey we observed this unique heterogenous habitat to be remarkably narrow with well-defined upper and lower limits. For this reason, it is conceivable that the Abyssinian Expedition simply “missed” sampling here along the survey route (nearest sampling localities in 1927: 3050 m and 3350 m). Note that in the likely event rising temperatures over the past century caused an upward shift in this transition zone, it would have been located lower in 1927 than it is today, and farther from the Abyssinian Expedition’s nearest sampled locality of 3350 m.

### Extreme small mammal endemism in SMNP

All 13 species documented by our survey are endemic to the Ethiopian Plateau (photographs of select taxa are provided in Supplementary Data SD4). Approximately half (54%; *n* = 7) are also endemic to the Simien Mountains, including three that resulted from type specimens collected during the Abyssinian Expedition and described by Osgood (i.e. *A. abyssinicus, C. baileyi*, and *L. simensis;* 1936). We expect to describe *C*. sp. indet. as a new taxon as well. Together, these four species account for well over half of our total collected specimens (61%; *n* = 289).

While all species are confined to Ethiopian highlands, some display more restricted ranges than others (Bryja et al. in press, for summary on rodents). Based on current taxonomic and biogeographic knowledge, we can separate rodents of SMNP into three main groups. The first comprises species relatively widespread in low to middle elevations of the Ethiopian highlands on both sides of the Great Rift Valley. In addition to *S. albipes*, we can also include the following in this group: *Dendromus mystacalis, Desmomys harringtoni*, and *M. mahomet*. The second group is formed by high-elevation species, endemic to the highest mountains in the north-western part of the Ethiopian highlands (*A. abyssinicus, L. simensis, O. simiensis, O. typus, S*. sp. “A”). Besides SMNP, most of these species are restricted to only a few other high mountain chains, e.g. Mt. Guna, Mt. Choqa or Abohoy Gara (Abuna Yosef). The third group is formed by *M. imberbis* and *D. lovati*—both species are known from the highest mountains on both sides of the Rift Valley (Bryja et al. in press; Meheretu et al. 2015), but are very rare or difficult to detect, and most records outside SMNP originate from Bale or Arsi Mts. in southeastern Ethiopia.

Both species of *Crocidura* are restricted to the highlands west of the Rift Valley, and SMNP is the only region from where they were confirmed genetically. *Crocidura baileyi* is considered a benchmark for the Simien Afroalpine community and Lavrenchenko et al. (2016) also mentioned its distribution in other mountains west of the Rift Valley (e.g. Debre Sina and Ankober). However, our recent genetic data suggest that high-elevation populations from Abohoy Gara Mts., Borena Saynt NP, and Ankober are very distinct—at least in mitochondrial DNA—and would require more taxonomic work. The second species, *C*. sp. indet., was first observed in a pitfall bucket at the 3600 m site and comprised 34 of the 123 (28%) soricids. It is a small, dark, shrew with an overall appearance that conspicuously differentiates it from other members of the genus in the region (total length, ca. 85 mm; tail length, ca. 35 mm; weight, ca. 3 g). Awaiting proper integrative taxonomic analysis, analysis of cytochrome *b* sequences suggests it may represent a sister taxon to *C. bottegi* (Lavrenchenko et al. 2009). In summary, SMNP can be considered an extremely valuable core area of Ethiopian endemism while also representing the best protected natural habitat in northern Ethiopia. Continued protection of SMNP will not only safeguard these rare and endemic small mammals, but also provide a potential refuge for lower elevation species responding to rising global temperatures.

### The role of elevation: diversity and abundance

Species richness was greatest at the 3250 m site, with eight rodent and two shrew species. This is consistent with a pattern of peak species richness for non-volant small mammals at mid-elevations as has been reported by several montane studies worldwide (Brown 2001; Goodman and Rasolonandrasana 2001; McCain 2004; Rickart et al. 2011; Stanley et al. 2014; Stanley and Kihaule 2016). In the context of continental Africa, however, Taylor et al. argues (2015) such hump-shaped distributions are the exception rather than the rule. Other surveys of non-volant small mammals on Eastern African mountains that are similar in scale to the Simiens have produced mixed results. On Africa’s highest mountain, Mt. Kilimanjaro, peak species richness (14 species) was recorded at the mid-elevation of 3000 m before plummeting at the higher 3500 m and 4000 m sites (four and six species, respectively; Stanley et al. 2014). A recent survey of Mt. Kenya (which coincided with our SMNP survey in September and October, 2015) provides elevational distribution data for two opposing slopes of the massif, Chogoria (southeast) and Sirimon (northwest). While mid-elevations produced peak species richness on both slopes, the Sirimon slope recorded an additional peak at its lowest elevation (Musila et al. 2019). The Rwenzori Mountains with its exceptionally speciose small mammal community represents perhaps the greatest contradiction to the hump-shaped distribution hypothesis in Eastern Africa. Here, species richness was found to decrease monotonically with each increase in sampling elevation (Kerbis Peterhans et al. 1998). Fittingly, the distribution that most resembled our results in SMNP was that of the Bale Mountains (located opposite the Simiens on the east side of the rift valley; see Fig. 1). Here, Yalden (1988) recorded peak species richness at 3200 m within a similarly shaped overall distribution to SMNP. Yalden’s description of the elevational profile of habitats in Bale is also remarkably similar to what we observed in SMNP. Perhaps most noteworthy is the mention of a hetergenous zone of transition where the forest ends and erica bush begins until reaching “a sharp upper treeline at 3250 m” (Yalden 1988). The concentration of species occuring at this elevation in the Simien and Bale Mountains may exemplify a theory put forth by Brown (2001) that species richness may be amplified at a given elevation when two conditions are met: 1) species having different habitat requirements overlap in their distributions; and 2) this occurs at the most productive point in the gradient (Brown 2001).

Given the relative size of the Simien massif, 3250 m represents the truest middle elevation from our survey, as our lowest site of 2900 m near the base camp of SMNP was fairly high compared to the surrounding plateau (ca. 2000 m). The 2900 m site also recorded a substantially lower sample success of 1.3% (Table 2). Park staff informed us that livestock no longer grazed in the area of natural Afromontane forest in which we placed a portion of our traps (33% of buckets and 50% of traplines), however, we believe our results may reflect the effects of the former disturbance. Future surveys incorporating an alternate locality within this elevational range would help confirm whether or not this is the case.

Although not initially included as one of our objectives for the study, we found a correlation between elevation and average weight among congeners. For each of the five genera represented by two species (*Crocidura*, *Dendromus*, *Mus*, *Otomys*, and *Stenocephalemys*), the highest average weight consistently belonged to that of the higher elevation species. This observation may be explained by Bergmann’s initial rule (i.e. *interspecific* variation among congeners) as applied to elevation (Bergmann 1847). While studies have investigated the merits of Bergmann’s Rule *intraspecifically* within mammals by latitude (Taylor et al. 2015), none have tested the prediction between closely related taxa as it relates to elevation. Be that as it may, additional research would be required to differentiate our results from coincidence.

### Methodological implications

The trap types and techniques used at each site were effective in sampling the small mammal communities along the elevational transect. Only the lowest site (2900 m) recorded a new species beyond the second day of trapping (Fig. 4). Therefore, we are confident that our survey offers a complete assessment of the rodent and shrew communities occurring at each site. Previous surveys in Eastern Africa have achieved similar success using the same sampling protocol (Stanley and Hutterer 2007; Stanley et al. 2014; Stanley and Kihaule 2016). Included among these is a survey of Mt. Kilimanjaro that—like our survey—accumulated all species for a site on the first day of trapping at the highest elevation (4000 m; Stanley et al. 2014). The lower habitat heterogeneity at such high elevations may allow for sampling entire communities with greater efficiency, however additional research would be required to confirm this theory. Nevertheless, the ability to collect reliable data on species richness and abundance in short order is virtually always in the practical interests of those conducting community assessments.

Our trapping results underscore the necessary role of pitfall buckets in thoroughly sampling non-volant small mammal communities. Pitfall buckets are often more effective at capturing the smallest mammals (weight ca. < 10 g) when compared to Sherman and snap trap varieties. For example, both *C*. sp. indet. and *D. mystacalis* (weight ca. 3 g and 9 g, respectively) were captured exclusively in buckets. However, deviations from this general association do occur and should be considered to avoid sampling biases. For example, despite *M. mahomet*’s relatively small size (weight, ca. 8.5 g), it was only captured by traps. Conversely, the majority of *D. lovati* (weight, ca. 18 g) captures were found in buckets. This species has been reported to be “not very common, at least in trapping yields” as well as having an upper range limit of 3550 m (Dieterlen 2005). However, *D. lovati* was not particularly uncommon in our survey (*n* = 17), even at 4000 m. Sampling bias caused by the absence or underutilization of pitfall buckets in previous surveys may account for their ‘rarity’.

Rain was rare during the survey (see Supplementary Data SD1). However, we found a significant positive correlation between rainfall and *C*. sp. indet. captures at the 2900 m site. For the two sample-nights following rain at the 3600 m site, the number of *C*. sp. indet. captures again increased, whereas *C. baileyi* captures remained unaffected, or decreased. The 4000 m site experienced the greatest amount of rainfall, yet it had no discernable effect on *C. baileyi* captures (*C*. sp indet. was absent from this site). Many studies have found a correlation between rainfall and sampling bias for shrews outside tropical Africa (McCay 1996; Ford et al. 2002), however given our results it would be interesting to see whether or not other similar events are specific to tiny shrews which may share a more ‘fragile’ ecology because of their size.

### Conclusion

This year (2019) marks the 50^th^ Anniversary of SMNP. Since its establishment, the park has faced constant pressure from human activity in the region, such as livestock grazing, wood harvesting, and military conflict. At the same time, the climate has warmed by 1.5° C (Jacob et al. 2017). As increasing rates of global warming continues, many lower elevation species may track suitable habitats as they shift to higher elevations. There is still much to be learned about how small mammal communities are responding to climate change, and montane ecosystems such as SMNP provide a practical means to study these biotic variations along elevational gradients over time. Only by understanding these systems can we develop effective conservation strategies to defend against any future ecological consequences of climate change in these areas of high endemism inherently prone to species loss. Therefore, as Ethiopia’s remarkable endowment of endemic mammalian biota continues to be described and documented, so does the call for continued research and conservation.

## Acknowledgments

Funding for this study came from the Field Museum of Natural History: Gantz Family Collections Center, Tanzania Research Fund; and the Council on Africa. We are appreciative to The Ethiopian Wildlife Conservation Authority (EWCA) and Simien Mountains National Park office for their cooperation and assistance in carrying out this study. Mekelle University provided essential logistical support and transportation. Genetic analysis of selected small mammals was performed by A. Bryjová within the Czech Science Foundation project no. 18-17398S. B. Besch created maps of Ethiopia and the study area. S. Goodman, A. Hailemariam, Y. Kidane, B. Kahsay, B. Gebru, A. Birrara, and K. Craig provided critical field and logistical support. L. Heaney and B. Marks offered invaluable guidance and editorial advice. M.A. Rogers, R. Banasiak and L. Smith were vital in processing the collection. A. Birara, B. Gebre, F. Welegebrial, G. Brhane, G. Wondaya, W. Hayelom, Y. Kidanu and 3^rd^ year postgraduate (summer) and undergraduate (regular) biology students joined in the field training in SMNP in 2015.

