## Supplementary Data SD1 for "Small terrestrial mammal distributions in Simien Mountains National Park, Ethiopia: A reassessment after 88 years"

*Supplementary Data SD1.*—Climatic data, including mean ± standard deviation, range, and sample size of days measured (number of days receiving rainfall in parentheses) for each site sampled between September and November 2015 in Simien Mountains National Park, Ethiopia.

| Elevation (m) | Daily Minimum Temperature (°C) | Daily Maximum Temperature (°C) | Daily Rainfall (mm) |
| --- | --- | --- | --- |
| 2900 | 7.5°±0.8 | 24.7°±0.7 | 1.1±2.2 |
|  | 6.7°-8.4° | 24.0°-25.4° | 0-5 |
|  | N=4 | N=3 | N=5 (2) |
| 3250 | 3.7°±0.8 | 19.1°±0.9 | 0±0 |
|  | 3.0°-5.0° | 17.8°-20.3° | 0 |
|  | N=6 | N=5 | N=7 |
| 3600 | 2.7°±1.2 | 17.6°±4.1 | 0.6±1.3 |
|  | 1.4°-4.3° | 10.6°-21.2° | 0-3 |
|  | N=5 | N=5 | N=5 (1) |
| 4000 | 0.6°±0.7 | 16.1°±5.8 | 3.7±9.0 |
|  | -0.2°-1.6° | 6.4°-20.5° | 0-22 |
|  | N=6 | N=5 | N=6 (1) |
