## Supplementary Data SD2 for "Small terrestrial mammal distributions in Simien Mountains National Park, Ethiopia: A reassessment after 88 years"

*Supplementary Data SD2.*—New sequences submitted to GenBank.

| INIT | Field # | CYTB-genus | CYTB-species | GenBank | FMNH # | Reference |
| --- | --- | --- | --- | --- | --- | --- |
| WTS | 11909 | Crocidura | baileyi | MN223650 | 229511 | This study |
| WTS | 11920 | Stenocephalemys | sp. nov. A | MF685493 | 229950 | Bryja et al. (2018) |
| WTS | 11922 | Stenocephalemys | sp. nov. A | MN223622 | 229952 | Bryja et al. (2018) |
| WTS | 11951 | Otomys | typus | MN223588 | 229707 | This study |
| WTS | 11952 | Dendromus | lovati | MN223635 | 229807 | This study |
| WTS | 11955 | Stenocephalemys | sp. nov. A | MF685494 | 229660 | Bryja et al. (2018) |
| WTS | 11957 | Crocidura | sp. nov. indet. | MN223663 | 229513 | This study |
| WTS | 11959 | Arvicanthis | abyssinicus | MN223599 | 229618 | Bryja et al. (2019) |
| WTS | 11969 | Dendromus | lovati | MN223638 | 229582 | This study |
| WTS | 11973 | Lophuromys | simensis | MN223625 | 229820 | This study |
| WTS | 11974 | Crocidura | sp. nov. indet. | MN223662 | 229521 | This study |
| WTS | 11977 | Crocidura | baileyi | MN223653 | 229756 | This study |
| WTS | 11979 | Mus | imberbis | MN223606 | 229648 | This study |
| WTS | 11981 | Otomys | typus | MN223586 | 229996 | This study |
| WTS | 11985 | Lophuromys | simensis | MN223631 | 229821 | This study |
| WTS | 11989 | Arvicanthis | abyssinicus | MN223598 | 229899 | Bryja et al. (2019) |
| WTS | 11990 | Crocidura | baileyi | MN223645 | 229524 | This study |
| WTS | 11995 | Stenocephalemys | sp. nov. A | MN223623 | 229663 | Bryja et al. (2018) |
| WTS | 11998 | Crocidura | sp. nov. indet. | MN223664 | 229528 | This study |
| WTS | 12002 | Crocidura | baileyi | MN223654 | 229532 | This study |
| WTS | 12008 | Crocidura | baileyi | MN223648 | 229535 | This study |
| WTS | 12010 | Crocidura | baileyi | MN223652 | 229537 | This study |
| WTS | 12018 | Lophuromys | simensis | MN223629 | 229823 | This study |
| WTS | 12030 | Crocidura | baileyi | MN223655 | 229539 | This study |
| WTS | 12033 | Dendromus | lovati | MN223639 | 229584 | This study |
| WTS | 12035 | Crocidura | baileyi | MN223656 | 229540 | This study |
| WTS | 12040 | Otomys | typus | MN223587 | 230002 | This study |
| WTS | 12049 | Lophuromys | simensis | MN223628 | 229604 | This study |
| WTS | 12054 | Arvicanthis | abyssinicus | MN223600 | 229629 | Bryja et al. (2019) |
| WTS | 12056 | Dendromus | lovati | MN223640 | 229585 | This study |
| WTS | 12064 | Otomys | typus | MN223589 | 230004 | This study |
| WTS | 12080 | Crocidura | baileyi | MN223649 | 229545 | This study |
| WTS | 12082 | Crocidura | baileyi | MN223657 | 229546 | This study |
| WTS | 12101 | Dendromus | lovati | MN223641 | 229587 | This study |
| WTS | 12103 | Stenocephalemys | sp. nov. A | MF685495 | 229962 | Bryja et al. (2018) |
| WTS | 12110 | Stenocephalemys | sp. nov. A | MF685496 | 229968 | Bryja et al. (2018) |
| WTS | 12112 | Stenocephalemys | sp. nov. A | MN223619 | 229969 | Bryja et al. (2018) |
| WTS | 12122 | Otomys | typus | MN223591 | 230008 | This study |
| WTS | 12126 | Otomys | typus | MN223592 | 230009 | This study |
| WTS | 12128 | Otomys | typus | MN223590 | 229718 | This study |
| WTS | 12137 | Stenocephalemys | sp. nov. A | MN223620 | 229973 | Bryja et al. (2018) |
| WTS | 12150 | Arvicanthis | abyssinicus | MN223604 | 229635 | Bryja et al. (2019) |
| WTS | 12166 | Stenocephalemys | sp. nov. A | MN223618 | 229686 | Bryja et al. (2018) |
| WTS | 12167 | Stenocephalemys | sp. nov. A | MN223621 | 229975 | Bryja et al. (2018) |
| WTS | 12172 | Dendromus | lovati | MN223642 | 229589 | This study |
| WTS | 12174 | Arvicanthis | abyssinicus | MN223601 | 229638 | Bryja et al. (2019) |
| WTS | 12184 | Arvicanthis | abyssinicus | MN223602 | 229642 | Bryja et al. (2019) |
| WTS | 12192 | Arvicanthis | abyssinicus | MN223603 | 229935 | Bryja et al. (2019) |
| WTS | 12193 | Crocidura | baileyi | MN223647 | 229558 | This study |
| WTS | 12199 | Stenocephalemys | sp. nov. A | MN223617 | 229647 | This study |
| WTS | 12207 | Crocidura | baileyi | MN223658 | 229560 | This study |
| WTS | 12213 | Dendromus | lovati | MN223634 | 229810 | This study |
| EWC | 12214 | Stenocephalemys | albipes | MN223613 | 230016 | Bryja et al. (2018) |
| EWC | 12216 | Crocidura | sp. nov. indet. | MN223666 | 229770 | This study |
| EWC | 12220 | Stenocephalemys | albipes | MF685476 | 229693 | Bryja et al. (2018) |
| EWC | 12226 | Crocidura | sp. nov. indet. | MN223661 | 229773 | This study |
| EWC | 12230 | Stenocephalemys | albipes | MN223615 | 229695 | Bryja et al. (2018) |
| EWC | 12232 | Lophuromys | simensis | MN223624 | 229830 | This study |
| EWC | 12242 | Stenocephalemys | albipes | MN223614 | 229982 | Bryja et al. (2018) |
| EWC | 12243 | Crocidura | sp. nov. indet. | MN223665 | 229775 | This study |
| EWC | 12246 | Lophuromys | simensis | MN223627 | 229831 | This study |
| EWC | 12248 | Desmomys | harringtoni | MN223596 | 229813 | This study |
| EWC | 12250 | Crocidura | sp. nov. indet. | MN223660 | 229777 | This study |
| EWC | 12251 | Lophuromys | simensis | MN223630 | 229832 | This study |
| EWC | 12278 | Crocidura | sp. nov. indet. | MN223667 | 229565 | This study |
| EWC | 12279 | Stenocephalemys | albipes | MN223612 | 229983 | Bryja et al. (2018) |
| EWC | 12294 | Dendromus | lovati | MN223636 | 229593 | This study |
| EWC | 12296 | Stenocephalemys | albipes | MN223616 | 229701 | Bryja et al. (2018) |
| EWC | 12301 | Lophuromys | simensis | MN223626 | 229857 | This study |
| EWC | 12302 | Lophuromys | simensis | MN223632 | 229858 | This study |
| EWC | 12305 | Otomys | simiensis | MN223593 | 229726 | This study |
| EWC | 12306 | Mus | imberbis | MN223607 | 229651 | This study |
| EWC | 12313 | Dendromus | mystacalis s.str. | MN223643 | 229594 | This study |
| EWC | 12320 | Stenocephalemys | albipes | MF685477 | 229703 | Bryja et al. (2018) |
| EWC | 12328 | Crocidura | baileyi | MN223646 | 229571 | This study |
| EWC | 12329 | Dendromus | lovati | MN223637 | 229595 | This study |
| EWC | 12335 | Desmomys | harringtoni | MN223597 | 229646 | This study |
| EWC | 12338 | Mus | imberbis | MN223608 | 229655 | This study |
| EWC | 12339 | Stenocephalemys | albipes | MF685478 | 229706 | Bryja et al. (2018) |
| EWC | 12341 | Otomys | simiensis | MN223595 | 229730 | This study |
| EWC | 12344 | Crocidura | baileyi | MN223651 | 229573 | This study |
| EWC | 12358 | Dendromus | lovati | MN223633 | 229599 | This study |
| EWC | 12361 | Crocidura | baileyi | MN223644 | 229574 | This study |
| EWC | 12364 | Mus | imberbis | MN223605 | 229734 | This study |
| EWC | 12367 | Crocidura | sp. nov. indet. | MN223659 | 229789 | This study |
| EWC | 12369 | Stenocephalemys | albipes | MN223611 | 229989 | Bryja et al. (2018) |
| EWC | 12380 | Mus | mahomet | MN223610 | 229656 | This study |
| EWC | 12389 | Otomys | simiensis | MN223594 | 230012 | This study |
| EWC | 12398 | Mus | mahomet | MN223609 | 229945 | This study |
