## Supplementary Data SD3 for "Small terrestrial mammal distributions in Simien Mountains National Park, Ethiopia: A reassessment after 88 years"

*Supplementary Data SD3.*—Product-moment correlation of cumulative sample nights and four capture parameters. * = P≤0.05; ** = P≤0.01; † = correlation not applicable; parentheses represent strong but not significant correlations.

| Cumulative sample-nights correlated with (across) | Number of individuals | Number of species | New species added | Cumulative species |
| --- | --- | --- | --- | --- |
| Total |  |  |  |  |
| traps (rodents only) | -0.993** | (-0.739) | -0.814* | (0.652) |
| traps (all captures) | -0.991** | (-0.693) | -0.816* | (0.652) |
| buckets (shrews only) | -0.870* | -0.800* | (-0.652) | † |
| buckets (all captures) | -0.875** | (-0.742) | -0.888** | 0.829* |
| traps and buckets combined (all captures) | -0.991** | (-0.684) | -0.793* | 0.763* |
| 2900 m |  |  |  |  |
| traps (rodents only) | (-0.693) | † | (-0.707) | † |
| traps (all captures) | (-0.693) | † | (-0.707) | † |
| buckets (shrews only) | -0.313 | (-0.707) | (-0.707) | † |
| buckets (all captures) | -0.213 | -0.224 | (-0.707) | † |
| traps and buckets combined (all captures) | (-0.534) | (-0.707) | -0.530 | (0.866) |
| 3250 m |  |  |  |  |
| traps (rodents only) | -0.817* | -0.476 | -0.791* | (0.612) |
| traps (all captures) | -0.813* | -0.455 | -0.757* | (0.612) |
| buckets (shrews only) | (-0.672) | -0.392 | (-0.612) | † |
| buckets (all captures) | -0.485 | -0.261 | -0.866* | 0.784* |
| traps and buckets combined (all captures) | -0.857* | -0.488 | -0.791* | (0.612) |
| 3600 m |  |  |  |  |
| traps (rodents only) | (-0.809) | -0.414 | -0.831* | (0.655) |
| traps (all captures) | -0.973** | -0.415 | -0.812* | (0.655) |
| buckets (shrews only) | -0.831* | (-0.703) | (-0.655) | † |
| buckets (all captures) | -0.855* | (-0.594) | (-0.655) | † |
| traps and buckets combined (all captures) | -0.973** | (-0.655) | (-0.724) | (0.655) |
| 4000 m |  |  |  |  |
| traps (rodents only) | -0.964** | -0.828* | (-0.655) | † |
| traps (all captures) | -0.967** | -0.828* | (-0.655) | † |
| buckets (shrews only) | (-0.588) | -0.207 | (-0.655) | † |
| buckets (all captures) | (-0.643) | -0.393 | -0.917* | 0.831* |
| traps and buckets combined (all captures) | -0.951** | -0.478 | (-0.655) | † |
