## Supplementary Data SD4 for "Small terrestrial mammal distributions in Simien Mountains National Park, Ethiopia: A reassessment after 88 years"

*Supplementary Data SD4.*—Field guide with representative photos of select small mammal species in Simien Mountains National Park, Ethiopia. Species 1-11 were sampled during our 2015 survey.

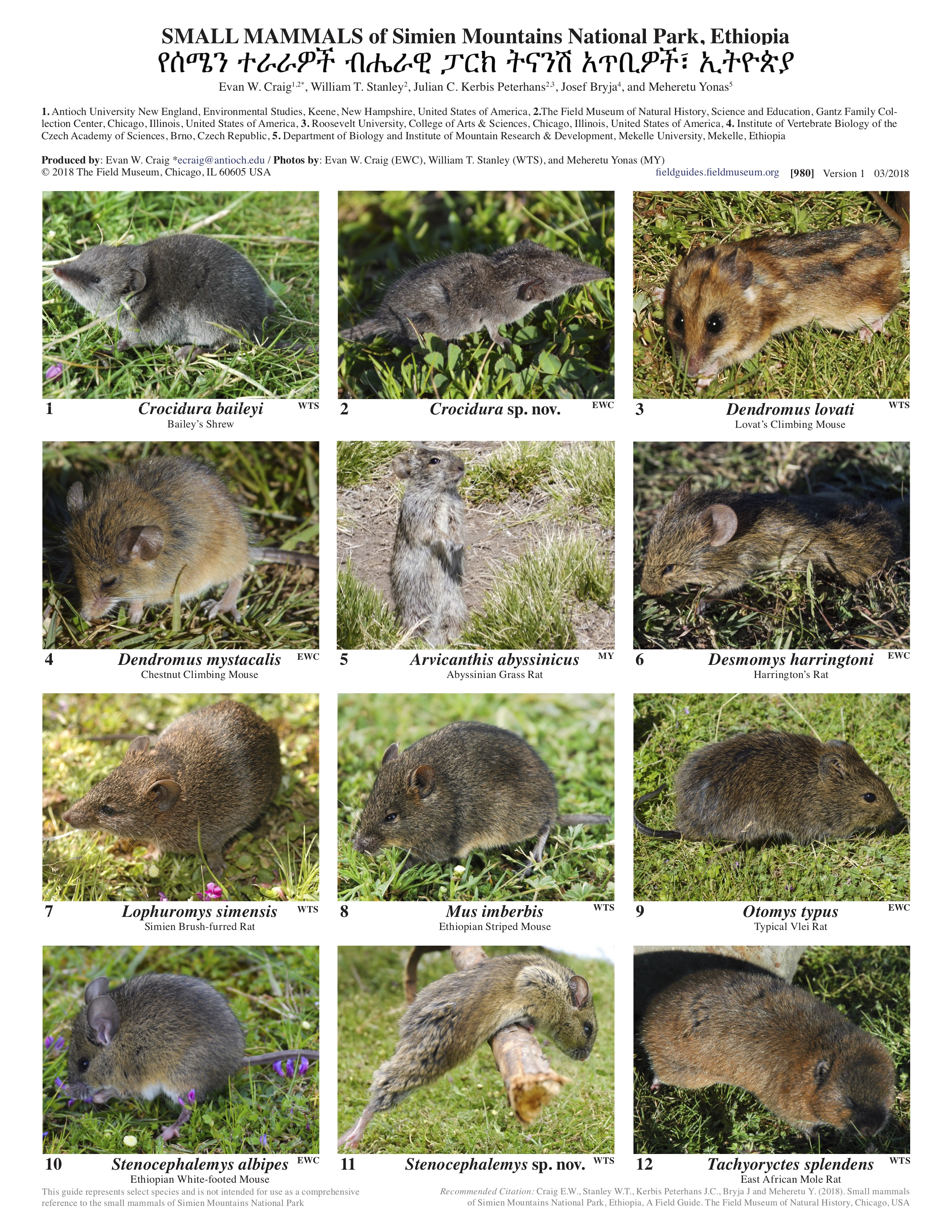
